# From zero to infinity: minimum to maximum diversity of the planet by spatio-parametric Rao’s quadratic entropy

**DOI:** 10.1101/2021.01.23.427872

**Authors:** Duccio Rocchini, Matteo Marcantonio, Daniele Da Re, Giovanni Bacaro, Enrico Feoli, Giles M. Foody, Reinhard Furrer, Ryan J. Harrigan, David Kleijn, Martina Iannacito, Jonathan Lenoir, Meixi Lin, Marco Malavasi, Elisa Marchetto, Rachel S. Meyer, Vítězslav Moudrý, Davnah Payne, Fabian D. Schneider, Petra Šímová, Andrew H. Thornhill, Elisa Thouverai, Saverio Vicario, Robert K. Wayne, Carlo Ricotta

**Affiliations:** BIOME Lab, Department of Biological, Geological and Environmental Sciences, Alma Mater Studiorum University of Bologna, via Irnerio 42, 40126, Bologna, Italy; Czech University of Life Sciences Prague, Faculty of Environmental Sciences, Department of Spatial Sciences, Kamýcka 129, Praha - Suchdol, 16500, Czech Republic; Department of Pathology, Microbiology, and Immunology, School of Veterinary Medicine, University of California, Davis, USA; Georges Lemaître Center for Earth and Climate Research, Earth and Life Institute, UCLouvain, Place Louis Pasteur 3, 1348 Louvain-la-Neuve, Belgium; Department of Life Sciences, University of Trieste, Via L. Giorgieri 10, 34127 Trieste, Italy; School of Geography, University of Nottingham, University Park, Nottingham NG7 2RD, UK; University of Zurich, Department of Mathematics and Department of Computational Science, Winterthurerstrasse 190, CH-8057 Zurich, Switzerland; Center for Tropical Research Institute of the Environment and Sustainability, University of California-Los Angeles, Los Angeles, CA 90095, USA; Plant Ecology and Nature Conservation Group, Wageningen University, Droevendaalsesteeg 3a, 6708 PB Wageningen, The Netherlands; Inria Bordeaux - Sud-Ouest, 200, avenue de la Vieille Tour, 33405 Talence, France; UR “Ecologie et dynamique des systèmes anthropisées” (EDYSAN, UMR 7058 CNRS-UPJV), Université de Picardie Jules Verne, 1 Rue des Louvels, 80037 Amiens Cedex 1, France; Department of Ecology and Evolutionary Biology, University of California- Los Angeles, Los Angeles, CA 90095, USA; Global Mountain Biodiversity Assessment, University of Bern, Institute of Plant Sciences, Altenbergrain 21, 3013 Bern, Switzerland; Jet Propulsion Laboratory, California Institute of Technology, 4800 Oak Grove Drive, Pasadena, CA 91009, USA; The University of Adelaide, Environment Institute, Adelaide, South Australia, 5000, Australia; State Herbarium of South Australia, Botanic Gardens and State Herbarium, Department for Environment and Water, GPO Box 1047, Adelaide, SA, 5001, Australia; CNR-IIA C/O Physics Department “M. Merlin” University of Bari, Via G. Amendola 173 - 70126 Bari, Italy; Department of Environmental Biology, University of Rome “La Sapienza”, Rome 00185, Italy

**Keywords:** biodiversity, ecological informatics, modelling, remote sensing, satellite imagery

## Abstract

**Aim:** The majority of work done to gather information on Earth diversity has been carried out by in-situ data, with known issues related to epistemology (e.g., species determination and taxonomy), spatial uncertainty, logistics (time and costs), among others. An alternative way to gather information about spatial ecosystem variability is the use of satellite remote sensing. It works as a powerful tool for attaining rapid and standardized information. Several metrics used to calculate remotely sensed diversity of ecosystems are based on Shannon’s Information Theory, namely on the differences in relative abundance of pixel reflectances in a certain area. Additional metrics like the Rao’s quadratic entropy allow the use of spectral distance beside abundance, but they are point descriptors of diversity, namely they can account only for a part of the whole diversity continuum. The aim of this paper is thus to generalize the Rao’s quadratic entropy by proposing its parameterization for the first time.

**Innovation:** The parametric Rao’s quadratic entropy, coded in R, i) allows to represent the whole continuum of potential diversity indices in one formula, and ii) starting from the Rao’s quadratic entropy, allows to explicitly make use of distances among pixel reflectance values, together with relative abundances.

**Main conclusions:** The proposed unifying measure is an integration between abundance- and distance-based algorithms to map the continuum of diversity given a satellite image at any spatial scale.

## 1 Introduction

Since Alexander von Humboldt (1769-1859), the spatial component of nature has played a relevant role in natural science. In the development of theoretical and empirical models in ecology, spatial structure represents a key concept to allow scientists to link ecological patterns to the generating processes and to the functional networking among organisms (Borcard and Legendre, 2002).

The majority of the work done to gather information about Earth diversity has been carried out by in-situ data, with known issues related to epistemology (e.g., species determination and taxonomy), spatial uncertainty, logistics (time and costs), among others (Rocchini et al., 2011).

Using satellite remote sensing can at least help attaining rapid and standardized information about Earth diversity (Gillespie, 2005; Rocchini et al., 2005). Furthermore, remote sensing can also be used to monitor some ecosystem functions and parameters such as temperatures, photosynthesis, vegetation biomass production and precipitation (Schimel et al., 2019; Zellweger et al., 2019) that can be useful to define the different niches of in-situ species, following first Goodall (1970) ideas, who envisaged future diversity measures as those based on niche theory (Hutchinson, 1959). The free access to remote sensing data (see Zellweger et al., 2019) has opened new ways to study ecosystem diversity and biodiversity issues (Rocchini et al., 2013). The spectral data related to pixels, as operational geographical units, are descriptions of pieces of land that allow us to define a new kind of Earth “diversity”, which may complement in-situ biodiversity measurement.

Diversity varies with area, thus investigating multiple spatial grains, until wide extents, is important to effectively monitor spatial diversity change in space and time (MacArthur et al., 1966). This is especially true in macroecology, where the primary aim is to model large-scale spatial patterns to infer the ecological processes which generated them, particularly considering the recent effect of global changes worldwide (Hobohm et al., 2019). In order to determine the horizontal distribution of diversity within a satellite image (i.e. which areas within the image are more diverse than others), diversity indices are usually spatially referenced by calculating the index within a moving window.

Several metrics that measure diversity from satellites rely on the Shannon’s theory of entropy (Shannon, 1948), with diversity being measured as 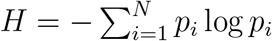, where *p*_*i*_ is the proportion of the *i*-th pixel value (e.g., digital number, DN) found within a moving window containing *N* pixels. Shannon’s *H* basically summarizes the partition of abundances (*sensu* Whit*taker, 1965*) by taking into account both relative abundance and richness of DNs (Figure 1).

**Figure 1:**
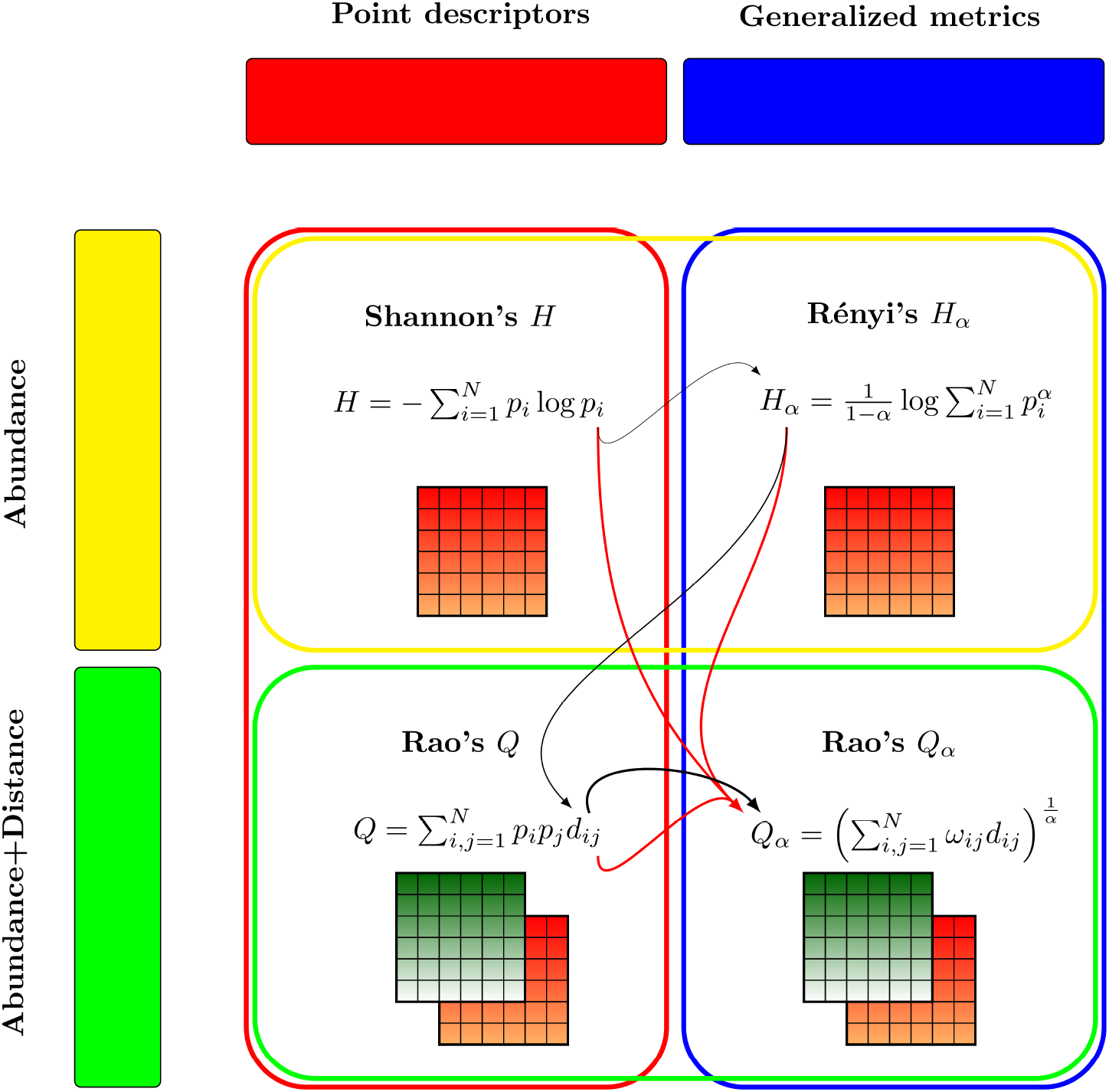
Grounding theory of this paper. Diversity measures can encompass abundance-based as well as abundance-distance-based metrics (yellow and green boxes, respectively). Abundance-distance-based metrics allow multiple layers to be used. The black lines represent the theoretical flow of this paper, with the thickness representing the complexity of each index, starting from Shannon’s Information Theory (point descriptor) to Rényi’s *H*_*α*_ (generalized entropy), which do not make use of distance. Distance enters the Rao’s *Q* formula, but this is still a point descriptor of diversity. Finally, parametric Rao’s *Q*_*α*_ comprises the use of distances and the generalized entropy concept. The red arrows represent the properties of the Rao’s *Q*_*α*_: i) it is grounded in Information Theory starting from Shannon’s H, ii) it is a generalized entropy like the Rényi *H*_*α*_, and iii) it makes use of distances like the Rao’s *Q*.

However, Shannon’s entropy is a point descriptor of (remotely sensed) diversity. As such, it shows only one part of the whole potential diversity spectrum at a glance. The use of generalized entropies has been advocated to face such problem. In this case, one single formula represents a parameterized version of a diversity index, thus providing a continuum of potential diversity indices. In the context of the measurement of diversity, the Rényi (1970) parametric entropy

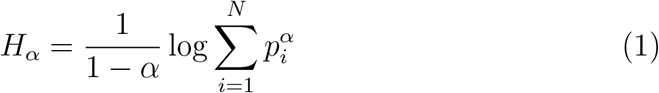

with 0 ≤ *α* ≤ ∞ represents a powerful tool to account for the continuum of diversity (Figure 1). One particularly convenient property of *H*_*α*_ is that by varying the parameter *α* there is a continuum of possible diversity measures, which differ in their sensitivity to rare and abundant DNs, becoming increasingly dominated by the most common DNs for increasing values of *α*. Note that for *α* →1, *H*_1_ equals the Shannon’s entropy. A similar formulation was then proposed by Hill (1973) who expressed parametric diversity as the “numbers equivalent” of Rényi generalized entropy. Appendix S1 provides the original formulation.

Rényi (and Hill) parametric functions summarize diversity by taking into account the pixel values of a satellite image and their relative abundances. However, they do not allow to explicitly consider the differences among these values. As an example, two arrays of 9 pixels with maximum richness and evenness (i.e. both containing 9 different DNs with relative abundances 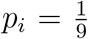) but differing in their values will attain the same Shannon diversity irrespective of the values of the DNs in both arrays.

By introducing a distance parameter *d*_*ij*_ among each pair of values *i* and *j*, Rao’s quadratic entropy (Rao, 1982)

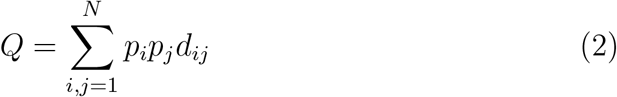

explicitly considers the differences among the pixel values in the calculation of diversity (Figure 1). Hence, two different pixels with values [2,3] will attain a lower diversity with respect to two pixels with values [0,100]. For instance, to make an ecological parallel, this is somewhat similar to the phylogenetic distance between two species: the values [2,3] would be equivalent to two sister species closely related on the tree of life while [1,100] would be equivalent to two very distant species on the tree of life.

The aim of this paper is thus to propose, for the first time, a parameterization of Rao’s quadratic entropy in order to provide a generalized entropy which accounts for both relative abundances and distances among pixel values. The proposed approach is now part of the `rasterdiv` R package, a package dedicated to diversity measures of spatial matrices, increasing its capability to discern among different diversity measures by a single formula.

## 2 Spatio-parametric Rao’s quadratic entropy

Inter-pixel spectral distances are directly related to landscape heterogeneity and they are capable of describing species habitats, starting with a satellite image (Rocchini et al., 2005). A satellite image can be viewed as a matrix of numbers describing Earth reflectance in different dimensions stored as pixels. A sensor per each light wavelength records the reflectance of a certain object in that wavelength which are stored into numbers in a certain range (e.g., digital numbers in 8 bits, ranging from 0 to 255). In general, the higher the variability in the spectral space defined by the pixel reflectance values, the higher the diversity of the ecosystem under study.

Consider a window of *N* pixels moving across the whole image to calculate a diversity index. Let *i* and *j* be two pixels randomly chosen with repetition within the moving window. Let *d*_*ij*_ be a symmetric measure of the (multi)spectral distance between *i* and *j* such that *d*_*ij*_ = *d*_*ji*_ and *d*_*ii*_ = 0. Rao’s *Q* (Rao, 1982) is defined as:

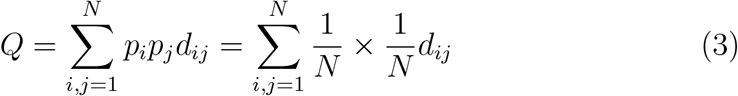

Therefore, *Q* measures the expected (i.e. mean) distance between two randomly chosen pixels and 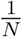 is the probability to extract each pixel. Note that, unlike *H*_*α*_ or *K*_*α*_ the calculation of Rao’s quadratic entropy is not limited to single bands but can be extended to multispectral systems of any dimension. For the connection between quadratic entropy and variance, see Rocchini et al., 2019.

Two parametric versions of quadratic entropy have been proposed by Ri-cotta and Szeidl (2006) and Leinster and Cobbold (2012). These parametric formulas were aimed at reconciling Rao’s *Q* with parametric entropies. However, they have only been rarely used in practice.

A more direct approach for developing a parametric version of quadratic entropy stems from the work of Guiasu and Guiasu (2011). Let 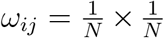 be the combined probability of selecting pixels *i* and *j* in this order. Guiasu and Guiasu (2011) noted that Rao’s *Q* can be expressed as a linear function of the combined probabilities of all pairs of pixels:

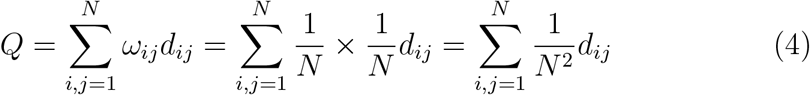

In practice, Rao’s *Q* is the arithmetic mean of the distances *d*_*ij*_ between all pairs of pixels *i* and *j*. Hence, in order to implement a parametric version of Rao’s *Q*, it seems natural to substitute the arithmetic mean in Equation 4 with a generalized mean (Hardy et al., 1952):

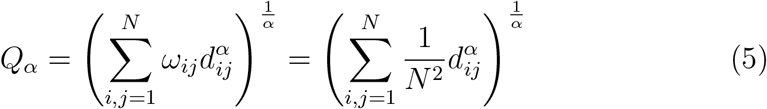

This operation connects *Q*_*α*_ with other diversity metrics that are expressed as generalized means, such as Hill’s (Hill, 1973) or Jost’s (Jost, 2006) numbers (Appendix S1) equivalents (see also Leinster and Cobbold, 2012).

The Rao’s *Q*, viewed as an arithmetic mean, is one of all the possible means in its generalized form *Q*_*α*_:

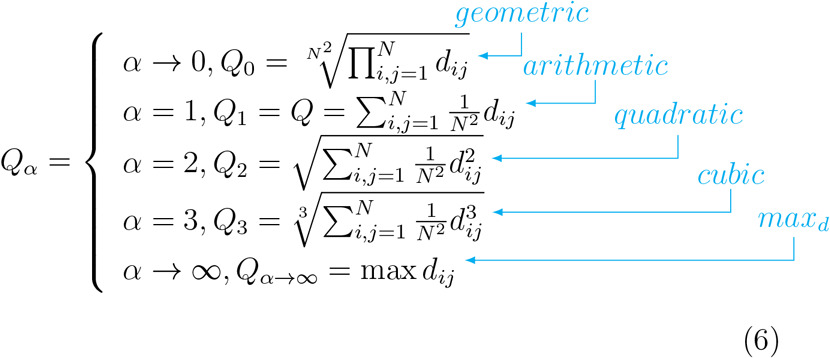

The mathematical proof that i) for *α* → 0 *Q*_0_ corresponds to the geometric mean, and ii) for *α*→ ∞ *Q*_∞_ corresponds to the maximum distance between pixel values pairs is provided in Appendix S1.

Each generalized mean always lies between the smallest and largest of its values. Increasing the parameter *α* will increase the weight of the highest values of *d*_*ij*_, thus providing a continuum of potential diversity indices (Figure 1).

## 3 The algorithm

Starting from a satellite image, a spatial moving window might be used to make the calculation on predefined extents of analysis. The grain (*sensu* Dungan et al., 2002) will be the resolution of the image while the extent of analysis will be the size of the moving window (Figure 2). The calculation is based on a distance matrix of type:

**Figure 2:**
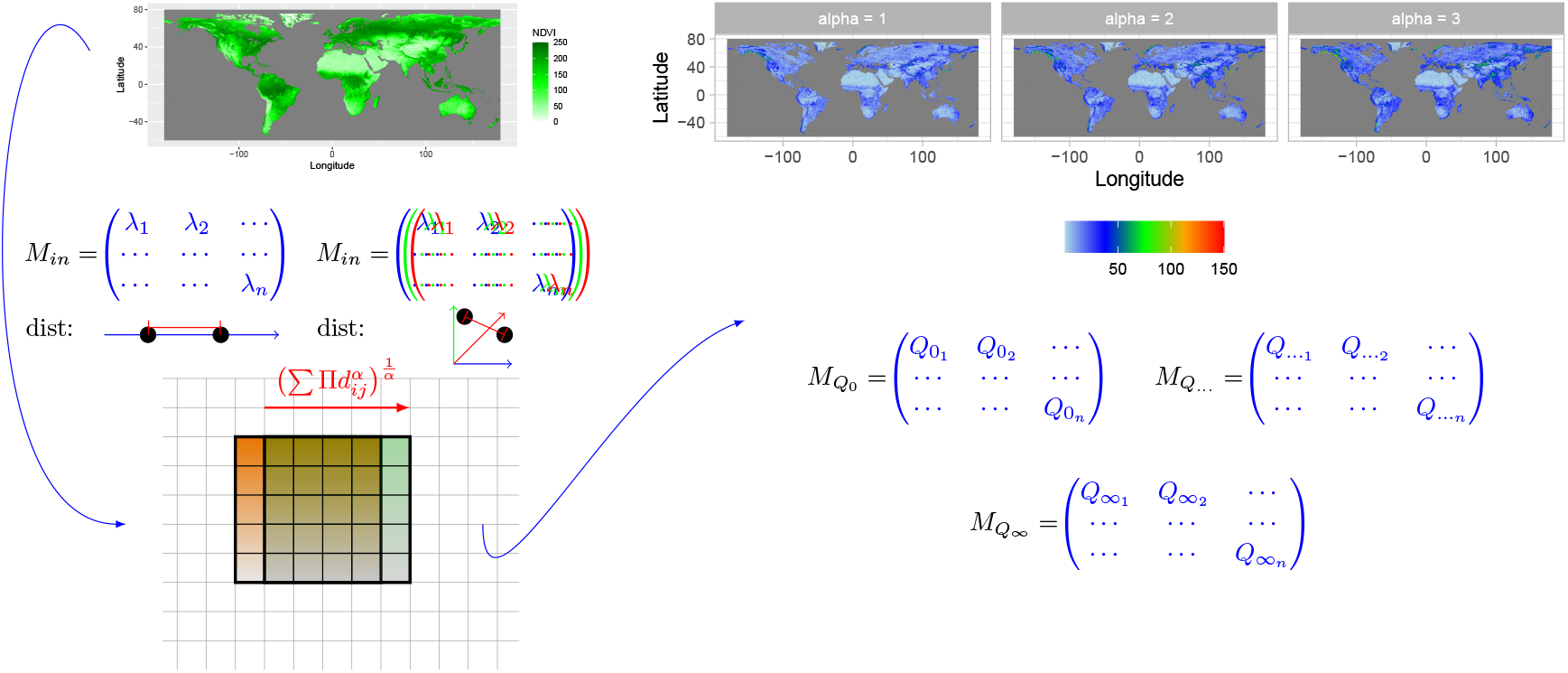
Starting from Copernicus Proba-V NDVI (Normalized Difference Vegetation Index) long term average image (June 21st 1999-2017) at 5km grain, parametric Rao’s *Q* is calculated in a moving window. In this paper NDVI was used as a single layer to calculate distances on one axis, but several layers can be used as well. In this example, three layers (blue, green and red matrices) are shown to calculate distances. The algorithm is based on a moving window passing throughout the whole image, calculating the Rao’s *Q*_*α*_ and saving the output in the central pixel. In this example a moving window of 5×5 pixels is passing (red arrow) from one position (orange) to the other (green). The output is a stack of layers each of which represents a different mean of the whole generalized mean spectrum of Equation 5.

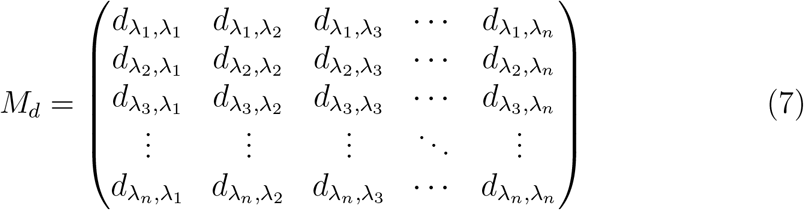

among all the potential pairs of pixels inside the moving window. The diagonal terms of the matrix (which equal zero) will have no effect for *α* > 0 (Equation 6), since they wou ld enter the term. On the contrary, for *α* → 0, they would enter the Π term by nullifying *Q*_0_.

We coded the proposed parameterization of Rao’s quadratic entropy as an R function, implementing the previously developed `rasterdiv` package (Mar-cantonio et al. (2020), https://CRAN.R-project.org/package=rasterdiv). The calculation of different *Q*_*α*_ by automatically changing the range of potential *α* values is done by the function `paRao`, as:

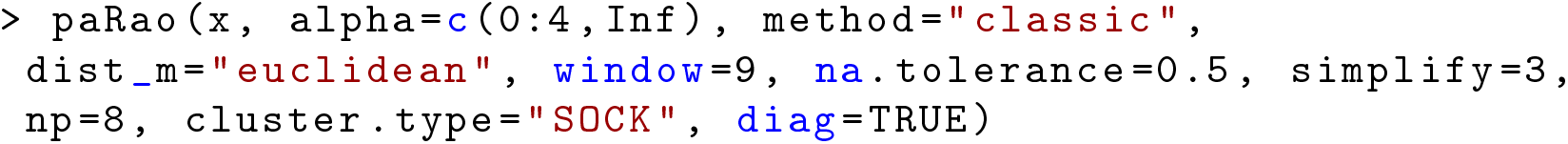

where x is the input dataset which can be a `RasterLayer` or a matrix class object, `alpha` is the *α* parameter of Equation 5, which can be a single value or a vector of integers. In the example above, *α* is a vector of integers ranging from 0 to 4, plus Inf, which in the R language is a reserved word representing positive infinity (*α*→ ∞). The option `method` decides if paRao is calculated with 1 single layer (`classic`) or with more than one layer (`multidimension`). With `method=“multidimension”` then x must be a list of objects. `dist m` is the type of distance considered in the calculation of the index, and can be set to any distance class implemented in the R package `proxy`, such as `“euclidean”`, `“canberra”` or `“manhattan”`. Moreover, `dist m` can also be an user-defined matrix of distances. However, if `method` is set to `“classic”` (unidimensional paRao) all distance types reduce to the Euclidean distance. The argument `window` is the side length in cells of the moving window (in this case set to 9), whereas `na.tolerance` is the proportion (0-1) of NA’s cell allowed in a moving window: if the proportion of NA’s cells in a moving window exceeds `na.tolerance` then the value of the moving window central pixel will be NA. The option `simplify` allows to reduce the number of decimal places to ease the calculation by reducing the number of numerical categories, i.e., if `simplify=3` only the first three digits of data will be considered for the calculation of the index. `np` is the number of parallel processes used in the calculation. If `np`>1 then the `doParallel` package will be called for parallel calculation, and `cluster.type` will indicate the type of cluster to be opened (default is `“SOCK”`, `“MPI”` and `“FORK”` are the alternatives). The `diag` argument refers to the diagonal term of Equation 7. It will have no effect on the function for *α* > 0, while it will nullify the value of *Q*_*α*_ if set to `TRUE`, as previously explained in Equation 7.

### 3.1 Global test of the parametric Rao’s Q variation over the planet

We applied the algorithm to a Copernicus Proba-V NDVI (Normalized Difference Vegetation Index) long term average image (June 21st 1999-2017) at 5km grain, also provided in the `rasterdiv` package as a free `Rasterlayer` dataset which can be loaded by the function `data()` (Figure 2). The parametric Rao algorithm can also be applied to multispectral data; in such a case distances are calculated in the multisystem created by the values of the pixels in each axis/band. The moving window passing throughout the whole image will return 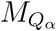 matrices/layers where *α* is the value chosen in the R function `paRao`.

With *α* → 0 the in Equation 6 leads to zeroes throughout the whole map (Figure 3). Increasing *α* will increase the weight of higher distances among different values until reaching the maximum distance value for *α*→∞. In this case the maximum turnover is reached and areas with maximum β-diversity will be apparent. In this case, a multitemporal set is used (long term average NDVI from June 21st 1999-2017). Hence, areas with the highest spatial and temporal turnover are enhanced, namely major mountain ridges. We expect that using single frame images would lead to the enhancement of the spatial component of diversity.

**Figure 3:**
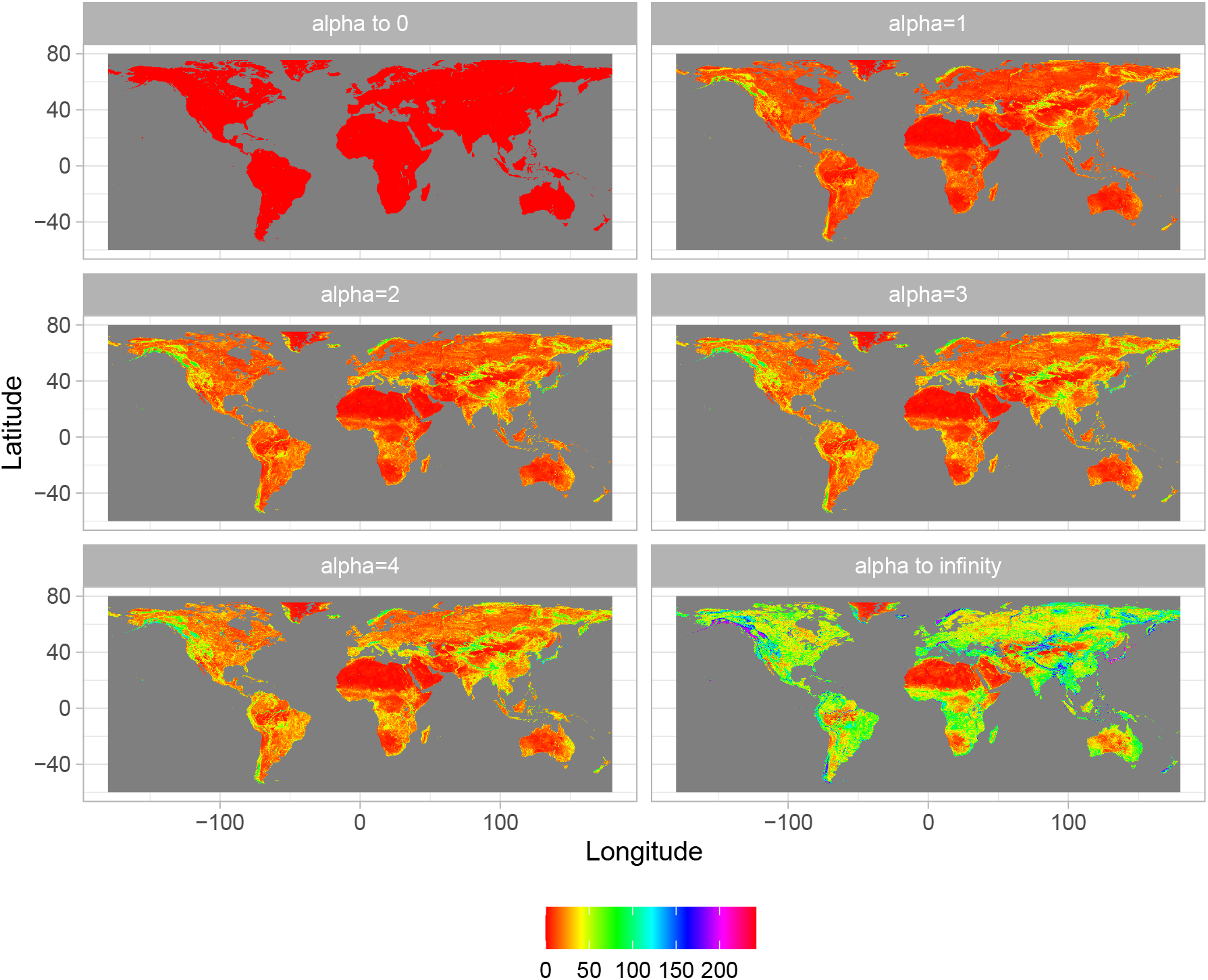
Output of the application of the algorithm shown in Figure 2, achieved by applying different *α* values: from 0 to 4 until *α* → ∞.. The higher the value of the parameter *α*, the higher the weight of highest distances among pixel values, until reaching the maximum potential *β*-diversity (maximum distance) at *α* → ∞.

Since the whole process is based on distances in a spectral space between pairs of pixels in terms of their “spectral characters” or in the “spectral space”, it is important to notice some cornerstone aspects on the use of distances from satellite images, especially when comparing different images or the same image in different times. In satellite images, the measure of distances could be impacted by: ii) the use of different sensors with different radiometric resolutions, as an example an 8-bit (2^8^ = 256 values) with respect to a 16-bit (2^16^ = 65536 values) image, or ii) the radiometric calibration which has been performed, e.g. with a non-linear transform. Therefore, care should be taken when making use of distances in remote sensing data, explicitly taking into account how the vector of proportions between pixels belonging to some defined classes (e.g., digital numbers, DNs) was obtained. The complete code of the function can be directly seen in R by typing the `paRao` function name. Moreover, a complete R coding session, to perform the above described analysis is provided in Appendix S2.

### 3.2 Local case study: the diversity of vegetation greenness and the ecoregions of California

A comparison between in-situ and remotely sensed diversity at worldwide scale might be difficult due to known biases in e.g. sampling effort, taxonomies, spatial uncertainty (Rocchini et al., 2017). Hence, we decided to calculate the Rao’s Q index on a NDVI raster layer of California (USA) to be compared with data in the field on native plant species diversity provided in Thornhill et al. (2017) from Baldwin et al. (2017). We chose California as a case study due to its high ecological diversity as well as to the availability of plant species field-data for this region.

In practice, we aimed at visualizing and describing differences in both diversity and structure of vegetation for the state of California, USA. First, an NDVI raster layer was derived from Copernicus Sentinel-2 data (European Space Agency, reference period: January 2017 to July 2018) and processed through Google Earth Engine to filter out cloud cover, select the greenest pixel of the time series and resample at 100 m pixel resolution. Then, the `paRao` R function was used to derive Rao’s Q index, considering both the original formulation of the Rao’s Q (*α* = 1, Equation 6) and the formulation with *α* → ∞ maximuzing *β*-diversity (Figure 3), with a moving window of 9×9 pixels.

A map of plant species richness was derived using the potential distribution range of 5,222 native California vascular plants modelled by Thornhill et al. (2017). Moreover, a vector map reporting the ecoregions of California (level III) was downloaded from the United States Environmental Protection Agency. In Figure 4, we showed NDVI, the Rao’s Q indices with *α* = 1 and *α*→ ∞ and plant species richness, reporting the boundaries of the different ecoregions for California. This comparison revealed macro-ecological and bio-geographical patterns which can be better interpreted considering the information condensed in the Rao’s Q index.

**Figure 4:**
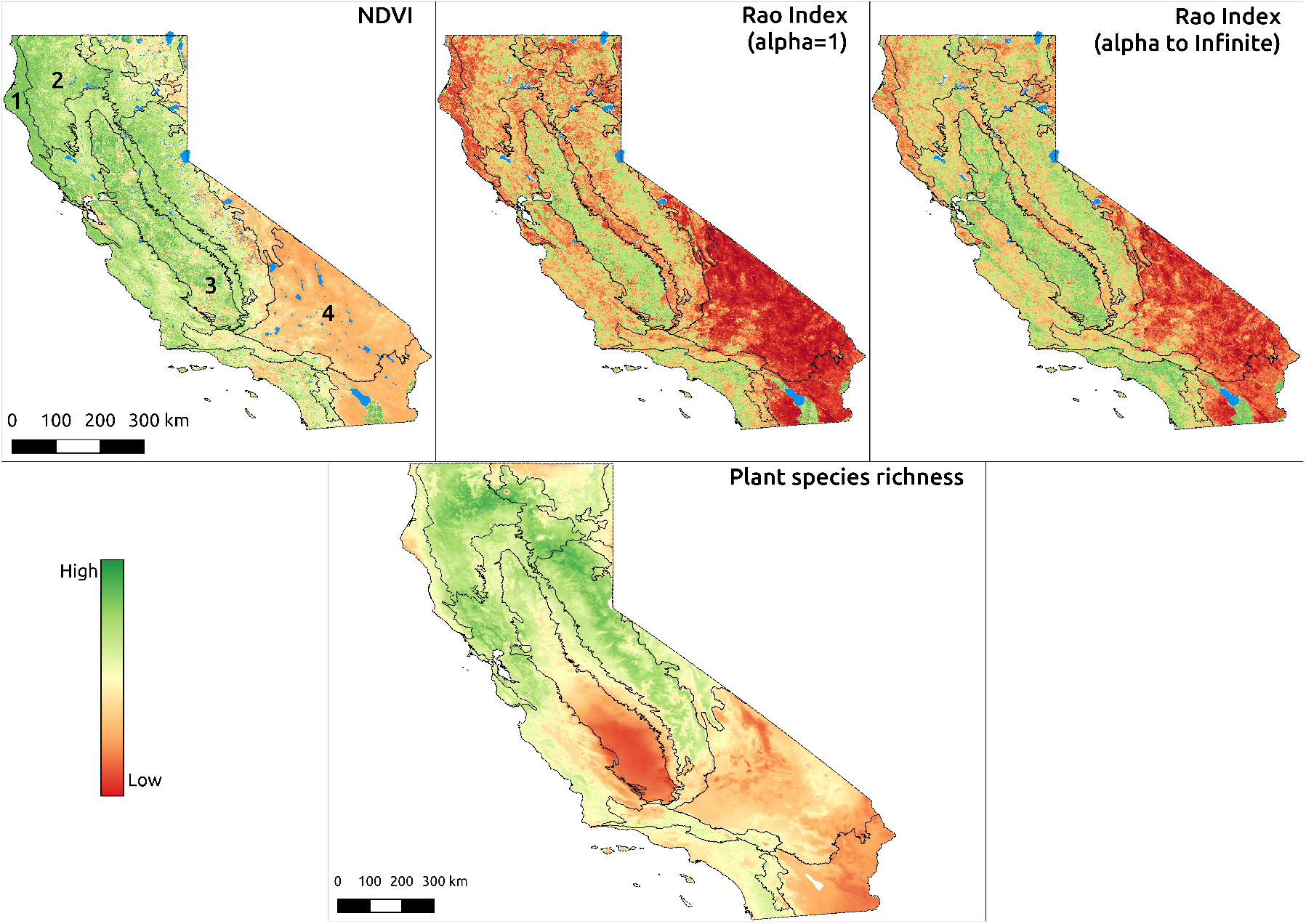
Comparison between NDVI, Rao’s Q Index, native plant species richness for the ecoregions of California. The NDVI values shown in the top-left box (100 m resolution) were derived from the ESA Copernicus Sentinel-2 dataset then processed with Google Earth Engine and range between -0.26 (red) and 0.99 (green). The Rao’s Q index shown in the top-right box was calculated from the NDVI map with alpha=1 and alpha to infinite and a moving window of 9×9 pixels. High values are shown in dark green and represent pixel whose sorrounding NDVI values are more “diverse” than pixel reported in red. The map reporting the potential native plant species richness of California (resolution: 810 m) was derived summing the binary potential distribution range of 5,222 native plant species modelled by Thornhill et al. (2017) and ranges between 134 (red) to 1029 (green) species per pixel (1 km^2^). The ecoregions considered in this paper are overlapped to the NDVI image: 1) Coast range (low mountains covered by highly productive, rain-drenched evergreen forests), 2) Klamath Mountain (highly dissected ridges, foothills, and valleys), 3) Central Valley (flat, urbanized and intensively farmed plains), 4) Mohave and Sonora ranges (very dry and warm broad basins).

For example, the ecoregion “Coast range” (labelled with 1 in Figure 4) is composed by low mountains covered by highly productive, rain-drenched evergreen forests. As a result, this region showed very high NDVI values but a low Rao’s Q index (low vegetation structural diversity) and low to medium plant species richness. The adjacent “Klamath Mountain” ecoregion (2) is instead characterized by highly dissected ridges, foothills, and valleys. This region still showed high NDVI values but higher Rao’s values with respect to region 1, which resulted in a high plant species richness. The diverse flora of this region, a mosaic of both northern Californian and Pacific Northwestern conifers and hardwoods, is rich in endemic and relic species. A similar pattern, although caused by opposite factors, was recognizable for the “Central Valley” region of California (3), which is composed of flat, urbanized and intensively farmed plains. The extensive presence of irrigated crops intersected with urbanized areas caused medium to high NDVI values and a very high apparent structural diversity. However, the same factors caused a low native species richness, especially in the drier southern portion of the valley. Finally, very dry and warm broad basins and scattered mountains characterize the “Mohave and Sonora ranges” ecoregions (4) which showed very low NDVI and Rao’s Q values (with scattered higher values associated with local topographical variability) and low native plant species richness.

Passing from the pure Rao’s Q index (*α*=1) to its parameterization with *α* → ∞ helped to increase the discrimination among areas, due to the fact that when *α*→ ∞ the Rao’s Q corresponds to the maximum distance (*β*-diversity) among pixel values in a site. Very similar gradients of the spatial heterogeneity of California (including BIOMOD variables, NDVI, elevation) as well as environmental DNA (eDNA) data are found in Lin et al. (2020).

## 4 Discussion

In this paper, we provided a straightforward solution to: i) account for distances in an Information Theory based metric, and ii) provide a generalized formula in order to avoid point description and account for the continuum of diversity. Diversity can be represented by different dimensions (Nakamura et al., 2020). Considering one single metric to account for the whole continuum of diversity metrics might be a powerful addition to the main framework. On the contrary, fragmenting the concept of diversity when trying to capture single aspects of the whole spectrum could be counterproductive.

The proposed unifying measure succeeded to integrate abundance- and distance-based algorithms over a wide variety of diversity metrics. We demonstrated that such integration is not only theoretical but also applicable to real spatial data, considering several dimensions of diversity at the same time. Being part of the `rasterdiv` R package, the proposed method is expected to ensure high robustness and reproducibility.

Remote sensing is obviously not a panacea for all the organismic based diversities like taxonomic-, functional-, genetic-diversity but it can represent an important exploratory tool to detect diversity hotspots and their changes in space and time at the ecosystem level. First of all, it measures heterogeneity of the environment with indirect links to the biodiversity of both plant and animal taxa, but also with potential discrepancies with species diversity, as in the presented case study of the native plant species diversity of California. This said, depending on the complexity and the resolution at which the proposed parameterized Rao’s *Q* is applied, it might allow finding new insights on the ecological processes acting in a certain ecosystem to shape its diversity. In this paper, the examples provided were based on a single NDVI layer since i) it is a valuable index of vegetation health and ii) it is freely available in the `rasterdiv` package to reproduce the code proposed in this paper. We are aware that NDVI has very limited capacity to track diversity in some habitats like dense forests, because it is saturated at dense vegetation. From this point of view, imaging spectroscopy offers higher information content, also enabling plant functional trait retrievals (Jetz et al., 2016; Schneider et al., 2019) as well as structural traits by LiDAR data (Schneider et al., 2020). The application of the proposed algorithm to future spaceborne imaging spectroscopy is promising. In other words, the algorithm has been thought to be used with multiple layers, like a whole multispectral image or the most meaningful Principal Components (Peres-Neto et al., 2005), or land use classes probabilities derived from fuzzy set theory (Rocchini and Ricotta, 2007; Feoli, 2018). This is even one of the major advantages of the Rao’s Q metric which allows considering both abundance and distance among pixel values, thus being applicable to any continuous raster layer, or to any matrix combination, even in a multiple spectral system.

Creating a unique “umbrella” under which all of the potential metrics of diversity can be used is highly beneficial for e.g. monitoring the variation in time of biological systems considering two major axes: i) the *α* parameter in Equation 5 providing information about the type of diversity at time *t*_0_, ii) the temporal dimension from time *t*_0_ to time *t*_*n*_ given the same *α* parameter. For the future, exploring such temporal dimension would allow gathering information of ecosystem changes in different diversity types at a glance.

Moreover, generalized entropy allows us to characterize the dimensionality of diversity (*sensu* Stevens and Tello, 2014) of different habitats/ecosystems. Those areas with a higher diversity dimensionality, namely a higher variability into the diversity spectrum would need a generalized measure to be fully undertaken. On the contrary, ecosystems with a lower dimensionality would have a lower difference among the different diversity measures with a flat curve of the diversity spectrum (Nakamura et al., 2020).

From a functional point of view, when all indices of diversity are highly correlated to each other (low dimensionality), it is expected that the ecological processes underlying diversity are just a few. On the contrary, with a lower correlation among indices (higher dimensionality) there might be a higher number of axes of variation coming out from different processes shaping ecological heterogeneity in space (Stevens and Tello, 2014).

There might be the possibility that a completely random matrix produces a pattern of diversity (Type I error). On the other side, a structured matrix could produce a very low diversity pattern (Type II error, Gotelli (2000)). In both cases, the parametric Rao’s *Q* could allow to determine, thanks to the use of a continuum of diversities, i) why a diversity pattern is still produced even in case of a random matrix, and ii) why a certain landscape shows a very low diversity in a certain point of the whole diversity spectrum. With point descriptors of diversity such inference cannot be done since the investigation is limited to a small window of the entire diversity spectrum, by basically relying on a single final number. In other words, the commonly asked question about what is the index which best describes diversity has no certain answer (Gorelick, 2011). Hence, the use of a trend of diversities will lead to the comprehension of hidden parts of the whole diversity dimensionality.

Furthermore, it is expected that the ecological processes shaping diversity should act at defined spatial scales (Borcard and Legendre, 2002). Hence, different diversity types of the whole dimensionality spectrum are expected to show scale dependent patterns, being apparent only at certain scales and not at some others. The use of a continuum allows measuring the different diversity types altogether in a single step. Changing the extent of analysis by different moving windows would then allow to encompass different spatial structures at different scales.

While geographic gradients of diversity over space are complex to catch in their very nature, biodiversity measurement has mainly relied in the past on few formulas which represented an hegemony (Stevens et al., 2013). In this paper, we demonstrated that diversity is actually multifaceted and should be necessarily approached through a generalized approach.

## 5 Conclusion

In order to unfold the dimensionality of diversity methods to directly account for several aspects of diversity at a time are needed. From this point of view, generalized entropy undoubtedly represents a powerful approach for mapping the diversity continuum.

Furthermore, it might be profitably used to plot multitemporal trends (see e.g. Dornelas et al., 2014) of diversity metrics and discover previously imperceptible differences when making use of single metrics (Figure 5).

**Figure 5:**
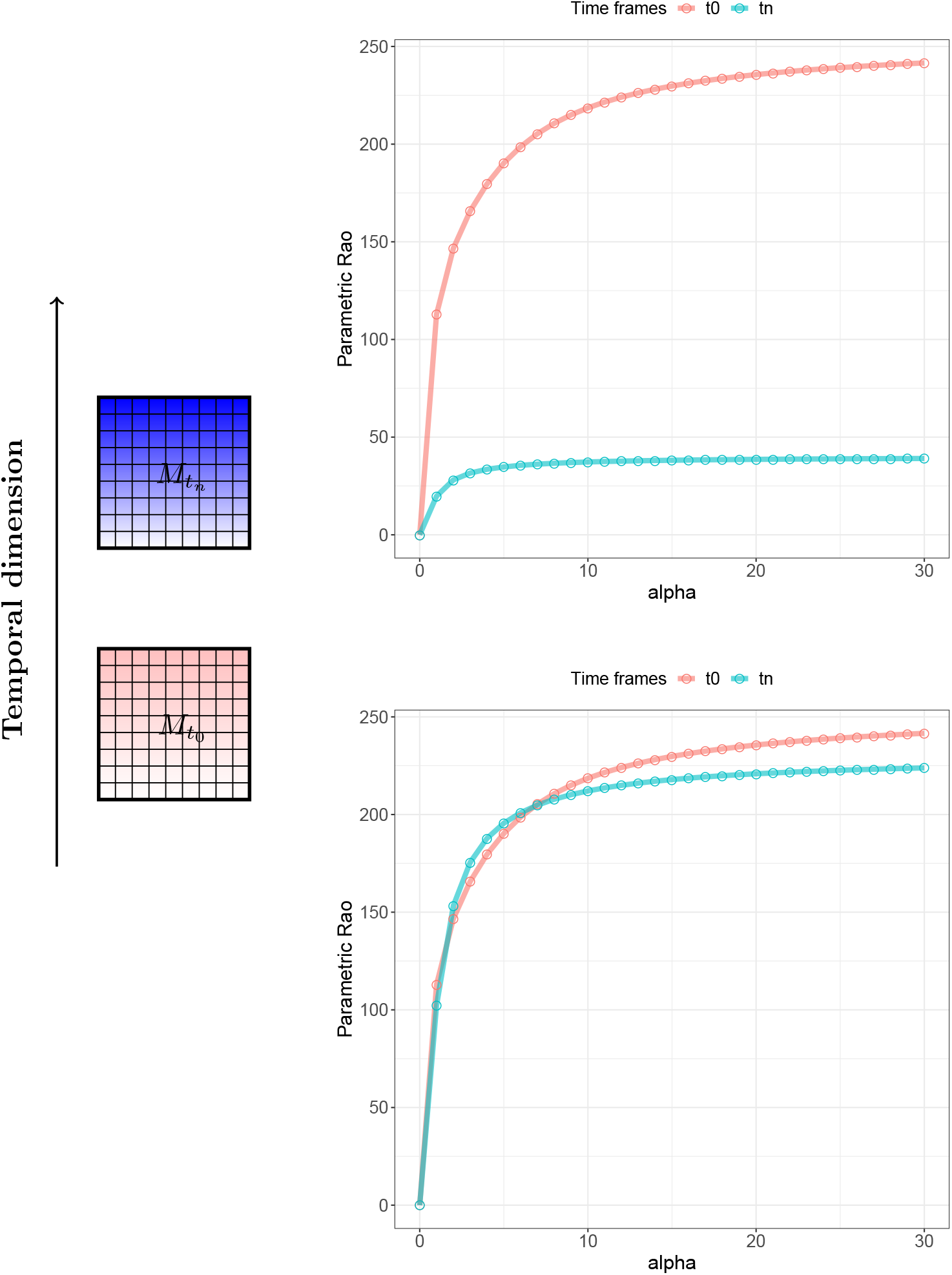
A theoretical example of the power of using generalized entropy for monitoring purposes. Given a landscape at times *t*_0_ (pink) and *t*_*n*_ (blue), calculating generalized entropy will allow the formation of a graph showing the continuum of Rao’s *Q* values observed over a range of values for *α*. The same landscape in different times might show an abrupt change (e.g., a catastrophic event) with an apparent diversity decrease (top). In this case, point descriptors (e.g., single *α* values) of diversity may be sufficient to describe this pattern. When the change in diversity is subtle (bottom), using a point descriptor might fail to detect it but it becomes manifest in the continuum of diversities based on generalized entropy. The complete code for reproducing this theoretical example is available in Appendix S3.

Metrics grounded in Information Theory ensure to make use of relative abundance of pixel values given the same richness in the moving window of analysis. However, distance metrics allow to also account for the relative dispersion in the spectral space of the cloud of pixels in a certain area (Laliberté et al., 2020). The proposed parameterization of the Rao’s *Q* explicitly considers the dispersion of pixel values in a spectral space (and their relative abundance) by allowing catching the whole dimensionality of diversity.

## 6 Data availability

The code and the data used in this paper are based on completely Free and Open Source Software, and they are available at the CRAN repository of the R package `rasterdiv`: https://CRAN.R-project.org/package=rasterdiv.

## 7 Acknowledgemnts

The research carried out at the Jet Propulsion Laboratory, California Institute of Technology, was under a contract with the National Aeronautics and Space Administration (80NM0018D0004). Government sponsorship is acknowledged. DR was partially supported by the H2020 project SHOWCASE and by the H2020 COST Action CA17134 “Optical synergies for spatiotemporal sensing of scalable ecophysiological traits (SENSECO)”.

## Appendix S1 Mathematical dissertation on the proposed algorithms

### 1 Hill’s numbers and generalized entropy

Hill (1973) expressed parametric diversity as the “numbers equivalent” of Rényi’s generalized entropy, as:

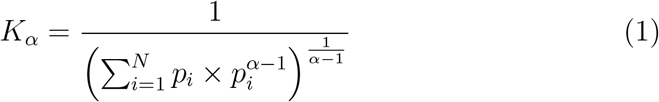

where the numbers equivalent *K*_*α*_ is the theoretical number of equally-abundant DNs (i.e. all those with 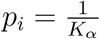) that are needed in order that its diversity be *H*_*α*_ (**?**).

Hill’s *K*_*α*_ has the form of the reciprocal of a generalized mean of order *α* − 1. Jost (2006) further showed that, like for *H*_*α*_, the numbers equivalents of all parametric and non-parametric measures of diversity that can be expressed as monotonic functions of 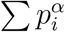 have the form of the reciprocal of a generalized mean of order *α* − 1 (for details, Jost, 2006).

### 2 Mathematical proof: for *α* → 0 *Q*_0_ is the geometric mean among the generalized means, for *α* → ∞ *Q*_∞_ is the maximum distance between pixel values pairs

We want to compute

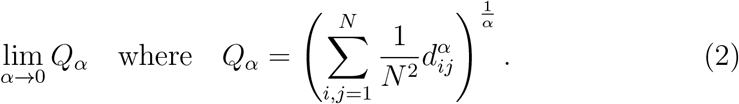

By exp(log(*x*)) = *x* we can rewrite *Q*_*α*_ as

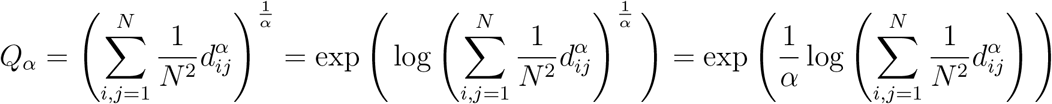

reminding that if *N* > 1, there is at least one distance *d*_*ij*_ > 0. We use this last expression to calculate (2). We use the following two well known results.

#### Theorem 1 (De l’Hôpital).

*Let f*_1_, *g*_1_: (*a, b*) ↦ ℝ *be two functions such that*

- lim_*x*→*a*_ *f*_1_(*x*) = lim_*x*→*a*_ *g*_1_(*x*) = 0
- *f*_1_ *and g*_1_ *are differentiable in* (*a, b*) *with* 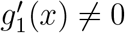 *for every x* ∈ (*a, b*)
- *the limit* 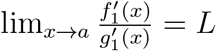 *with L* ∈ ℝ

*then*

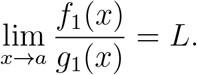

#### Theorem 2 (Limit composition).

*Let f*_2_: (*a, b*) ↦ ℝ *and let g*_2_: (*c, d*) ↦ ℝ *be two functions such that the image set of g*_2_ *is contained in the domain of f*_2_, *i*.*e*. ℐ*mg*(*g*_2_) ⊆ (*a, b*). *Let x*_0_ ∈ (*c, d*), *if it holds that*

- 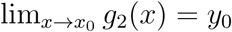 *with g*_2_(*x*) ≠ *y*_0_ *definitely for x* → *x*_0_
- 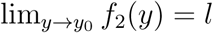

*with a, b, c, d, x*_0_, *y*_0_, *l* ∈ ℝ ∪ ±∞ *then*

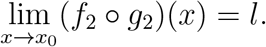

We apply Theorem (2) to calculate the limit (2) with *f*_2_(*x*) = exp(*x*) and

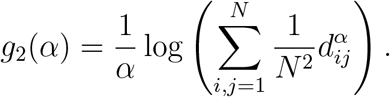

(all assumptions of the theorem hold). Setting *x*_0_ = 0, we have to compute

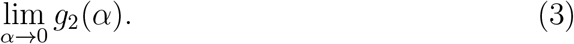

which will be accomplished using Theorem (1) by setting *f*_1_: (0, +∞) → ℝ

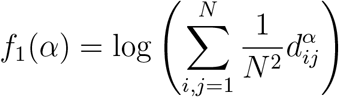

and *g*_2_: (0, +∞) → ℝ, *g*_2_(*α*) = *α*. Then we have

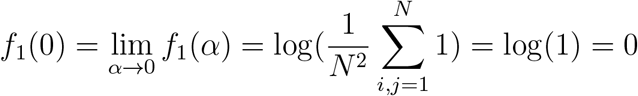

as the limit exists and

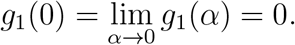

Both functions *f*_1_ and *g*_1_ are differentiable. Lastly we observe that 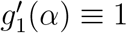. Since all the assumptions of Theorem 1 hold then

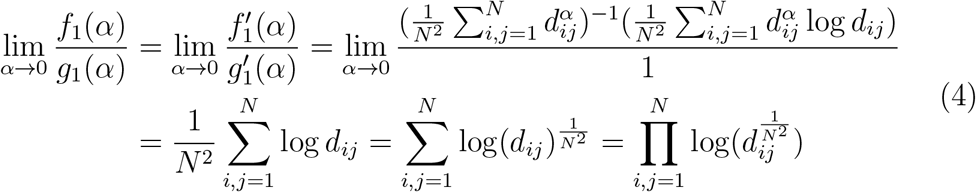

By Equation (4) we have the expression of Equation 3. Let

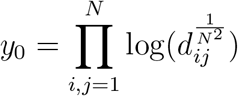

and we conclude by observing

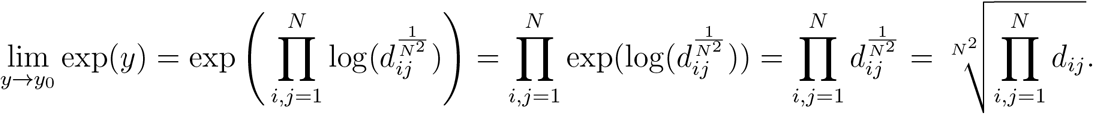

Now we want to compute

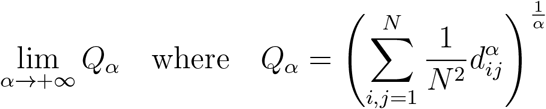

We define *d* = max{*d*_*ij*|_*i, j* ∈ {1, …, *N*}} and we rewrite *Q*_*α*_ as

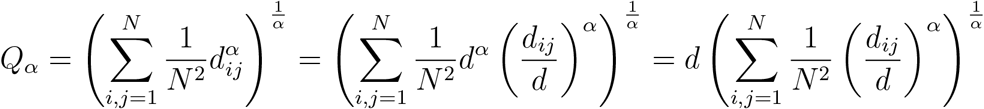

Next we observe that

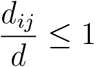

by construction and there exist a pair 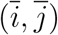 such that 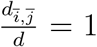. Therefore it follows that

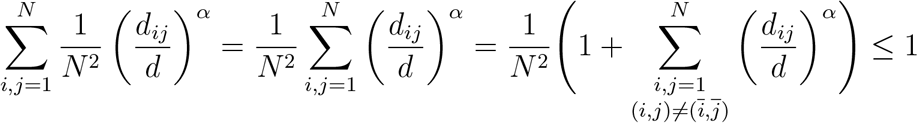

for every *α* > 1. And the limit in (4) is

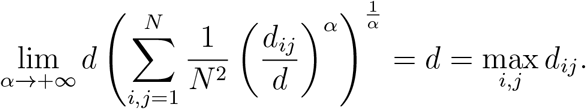

## Appendix S2 Code

### 1 paRao function

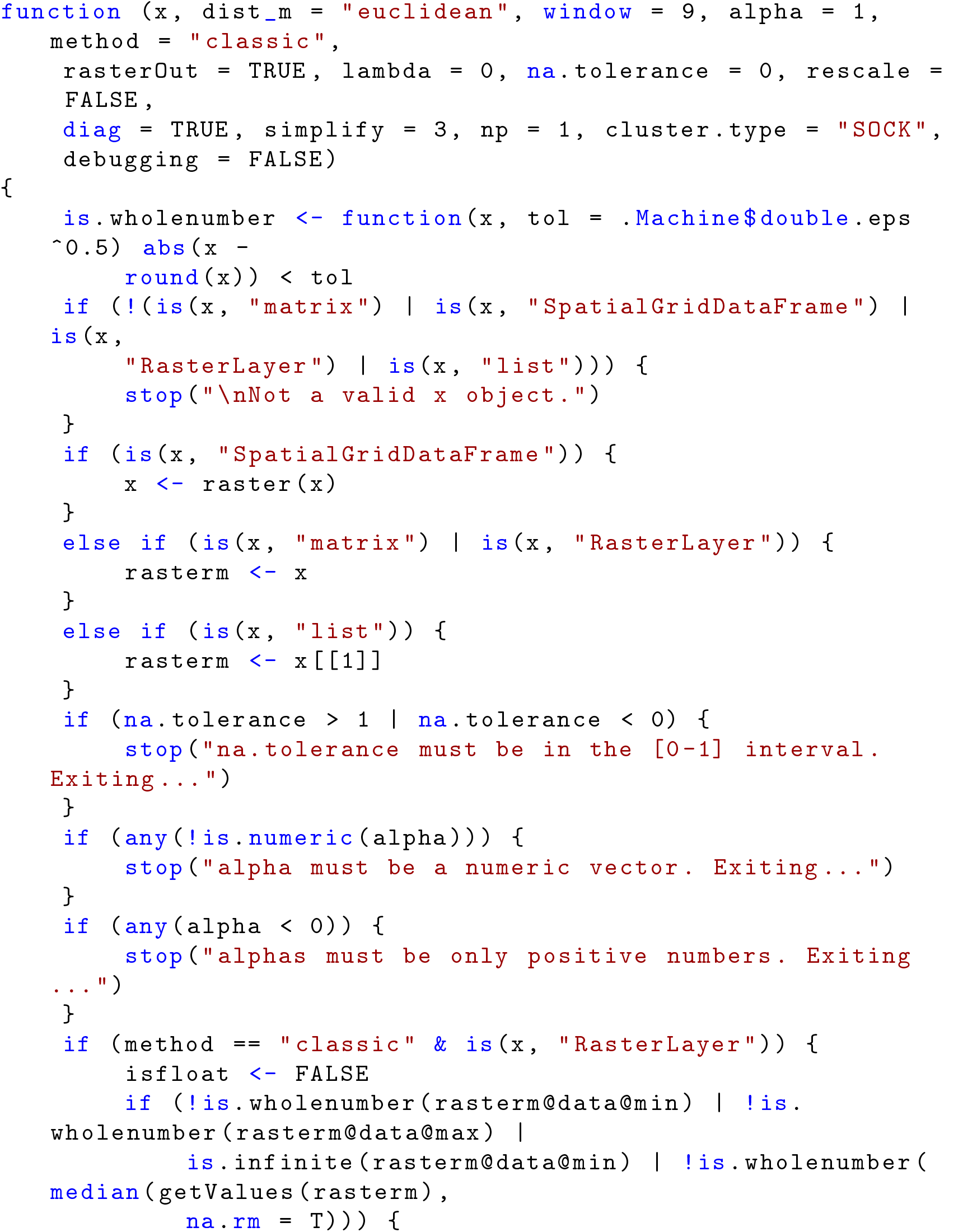
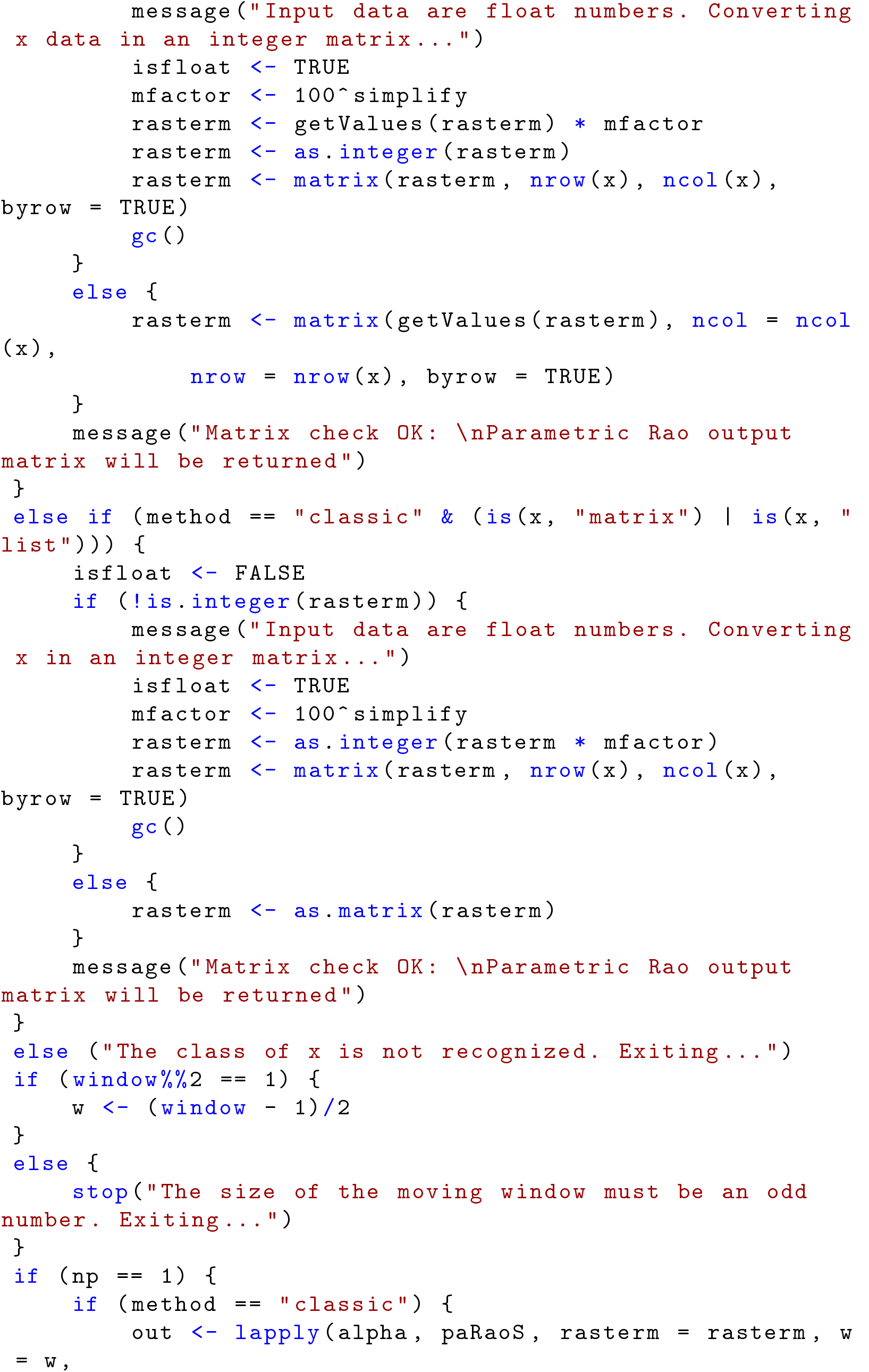
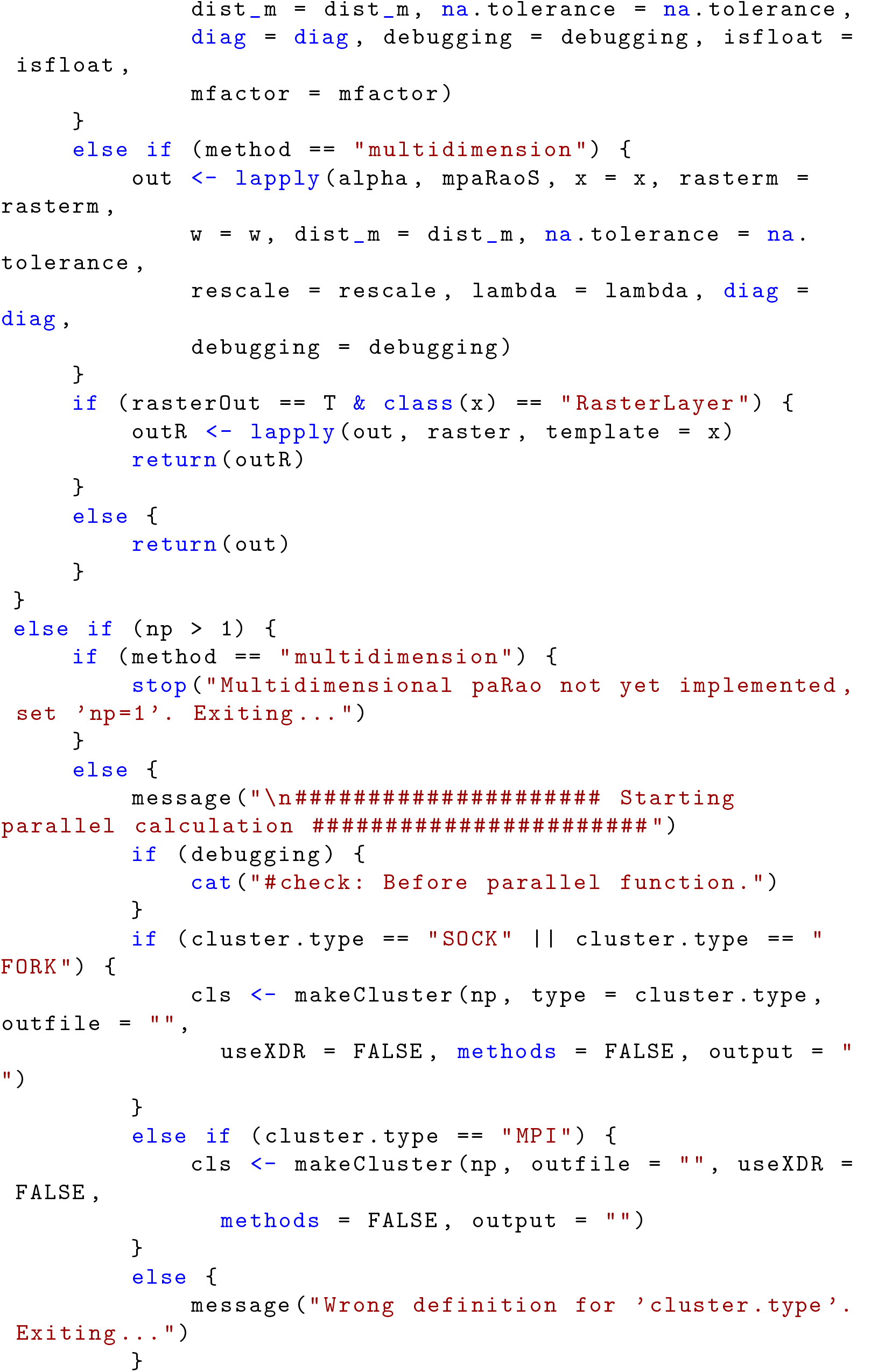
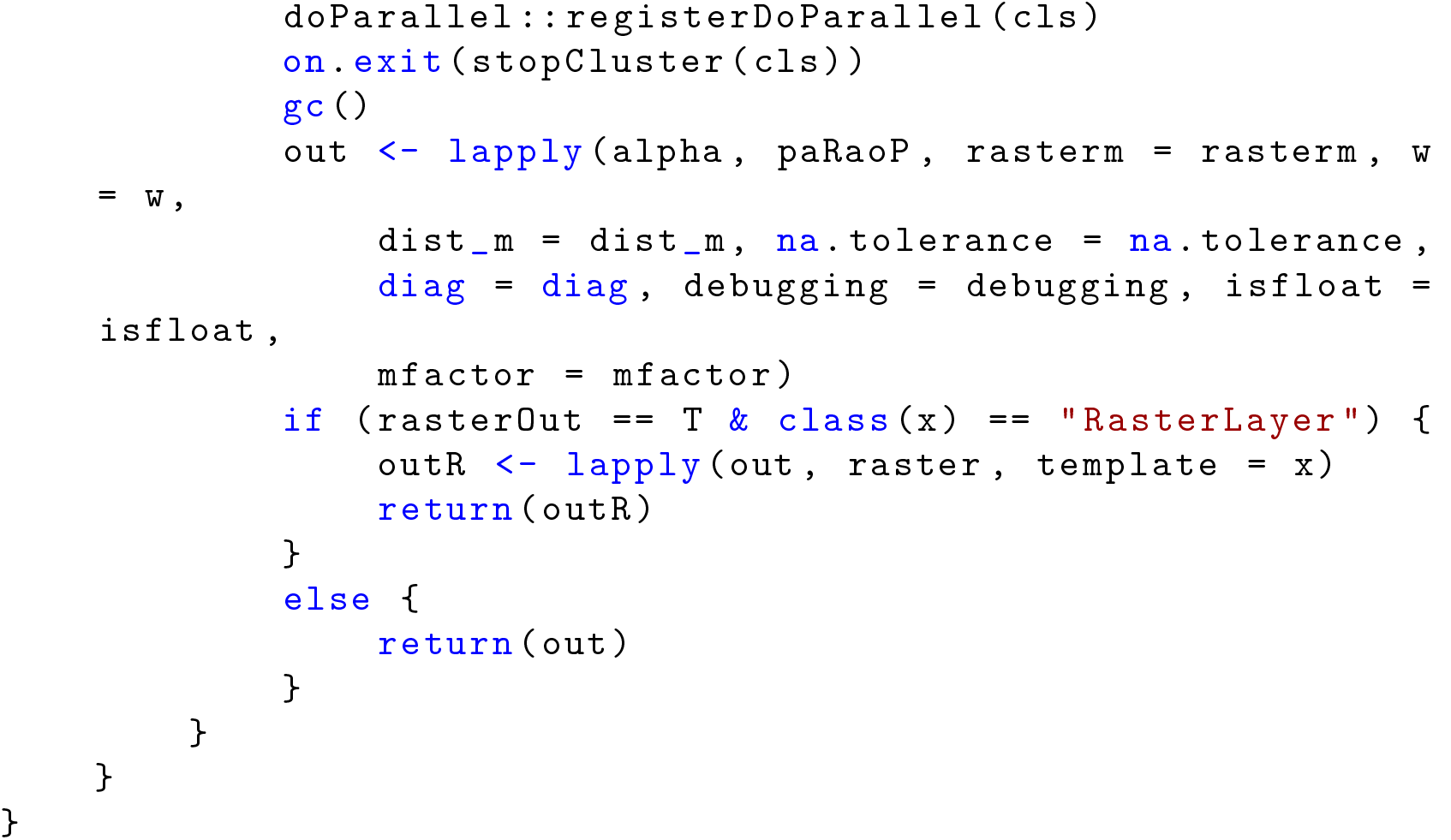

### 2 Application of the paRao function to a synthetic set

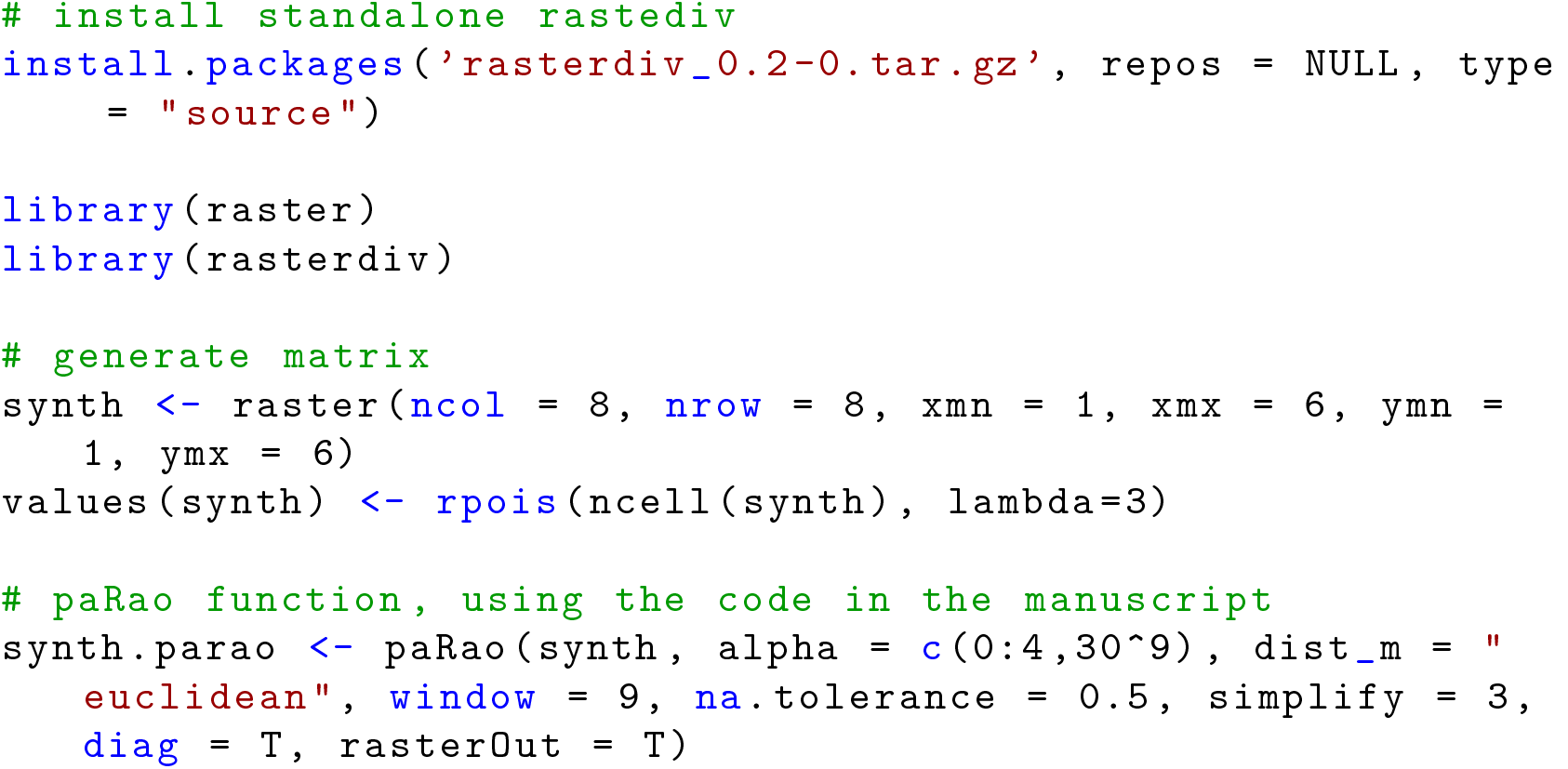

### 3 Application of the paRao function to the 8bit copNDVI dataset

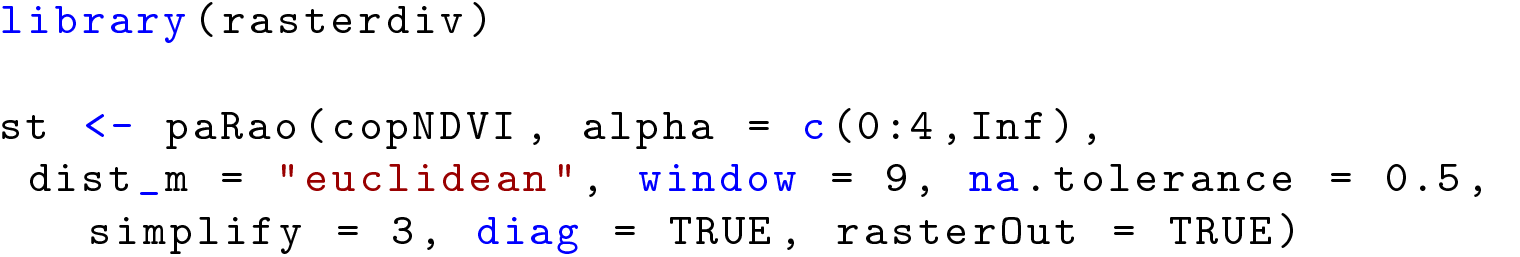

### 4 Output plot

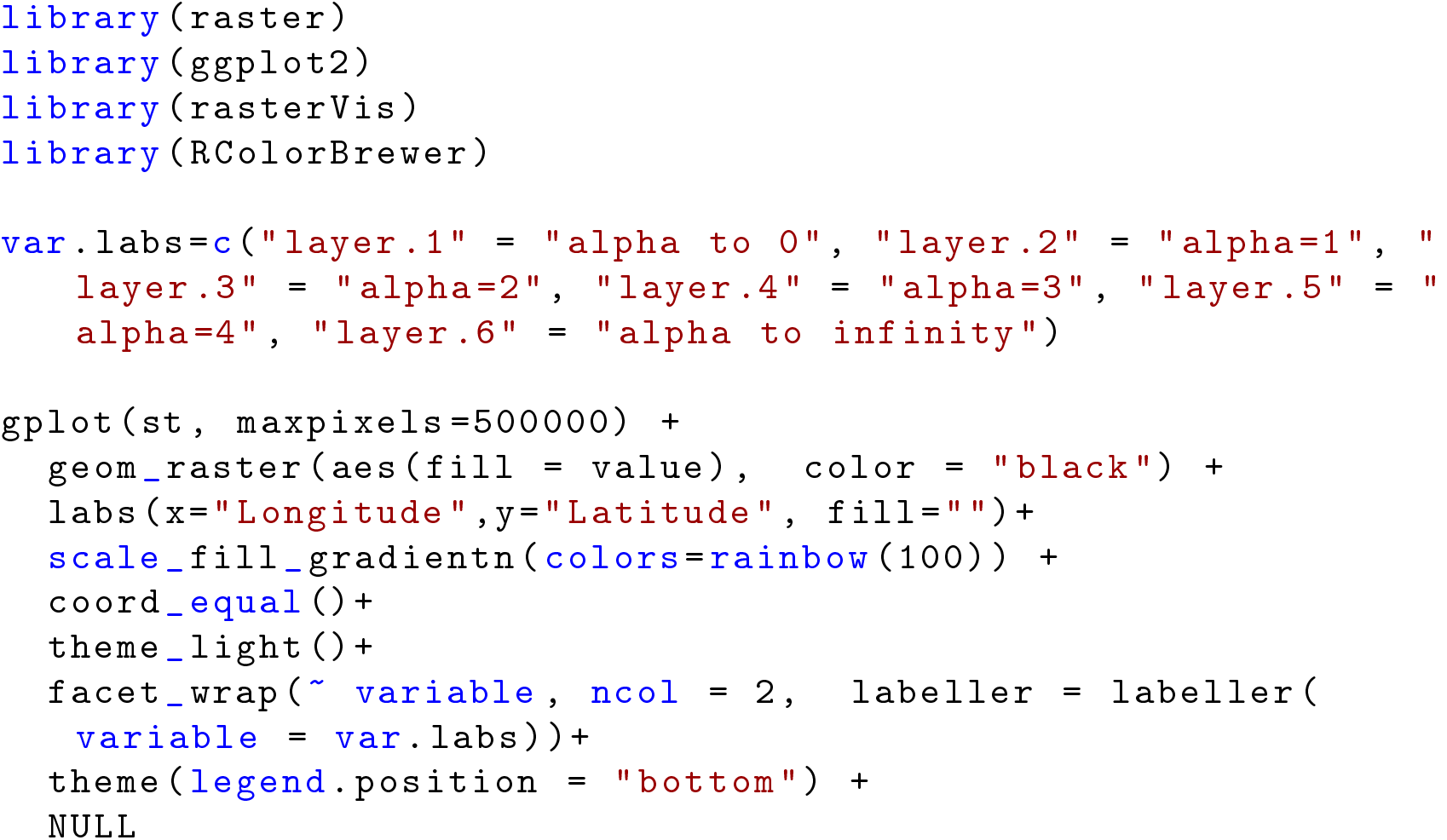

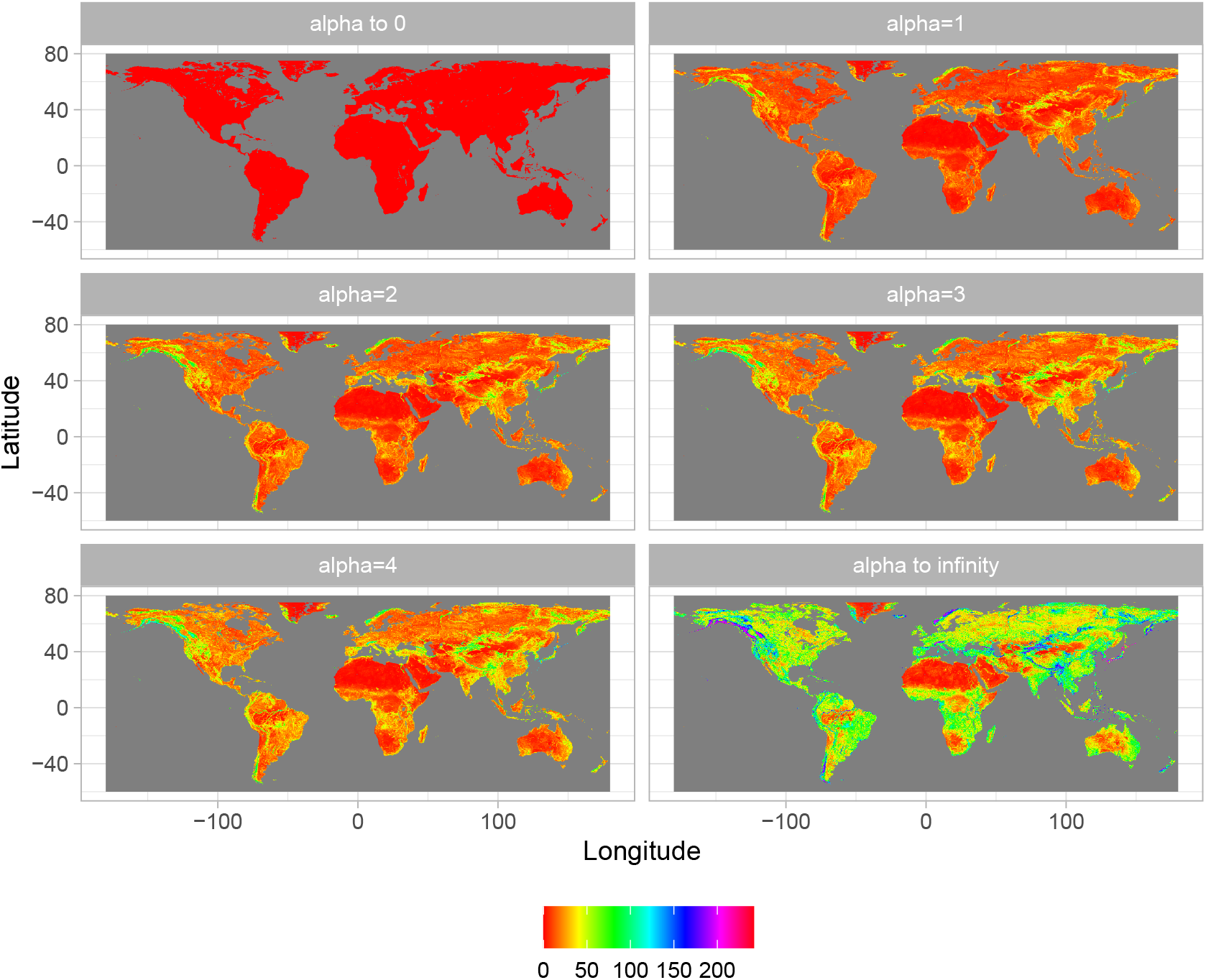

## Appendix S3 Code for Figure 4

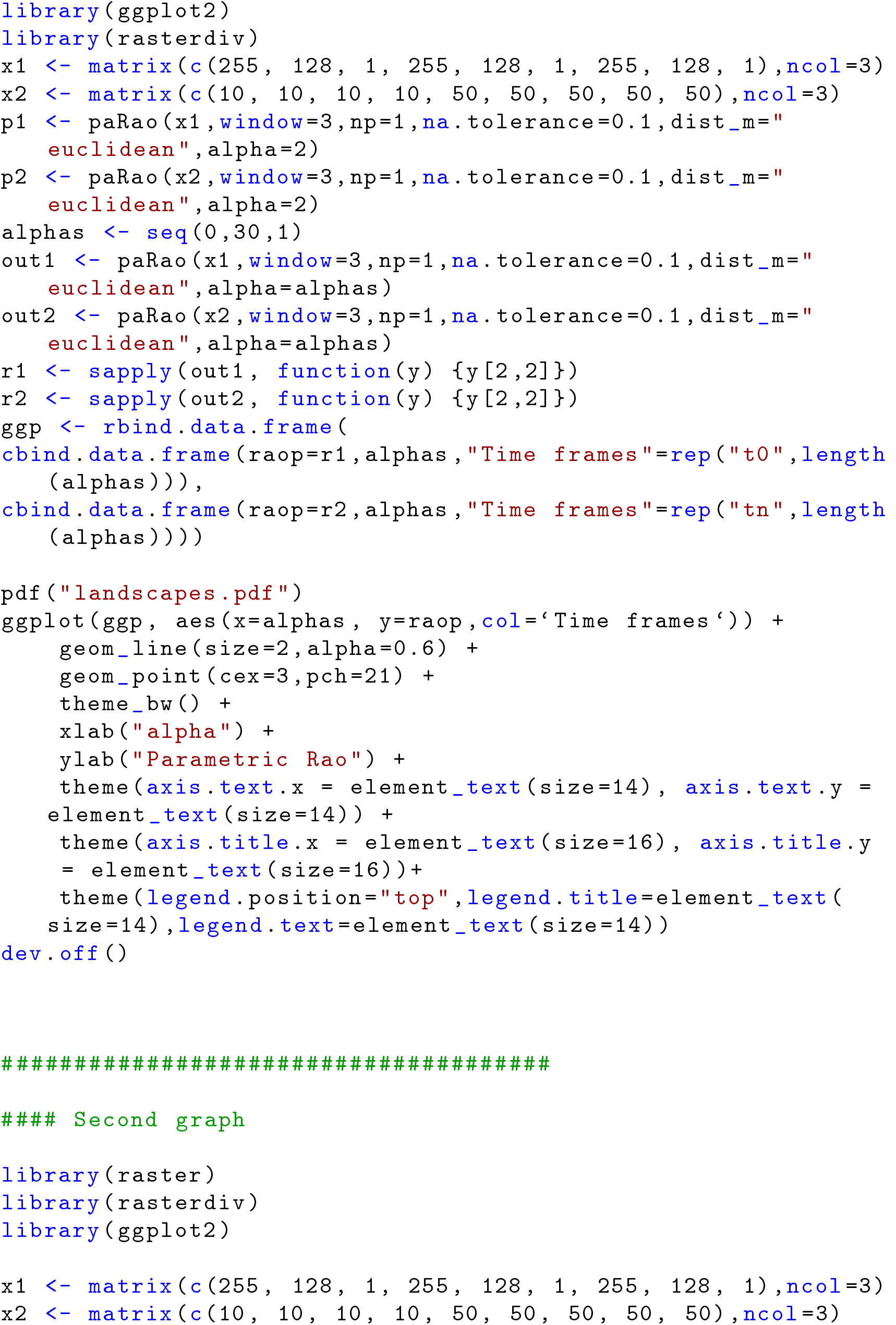
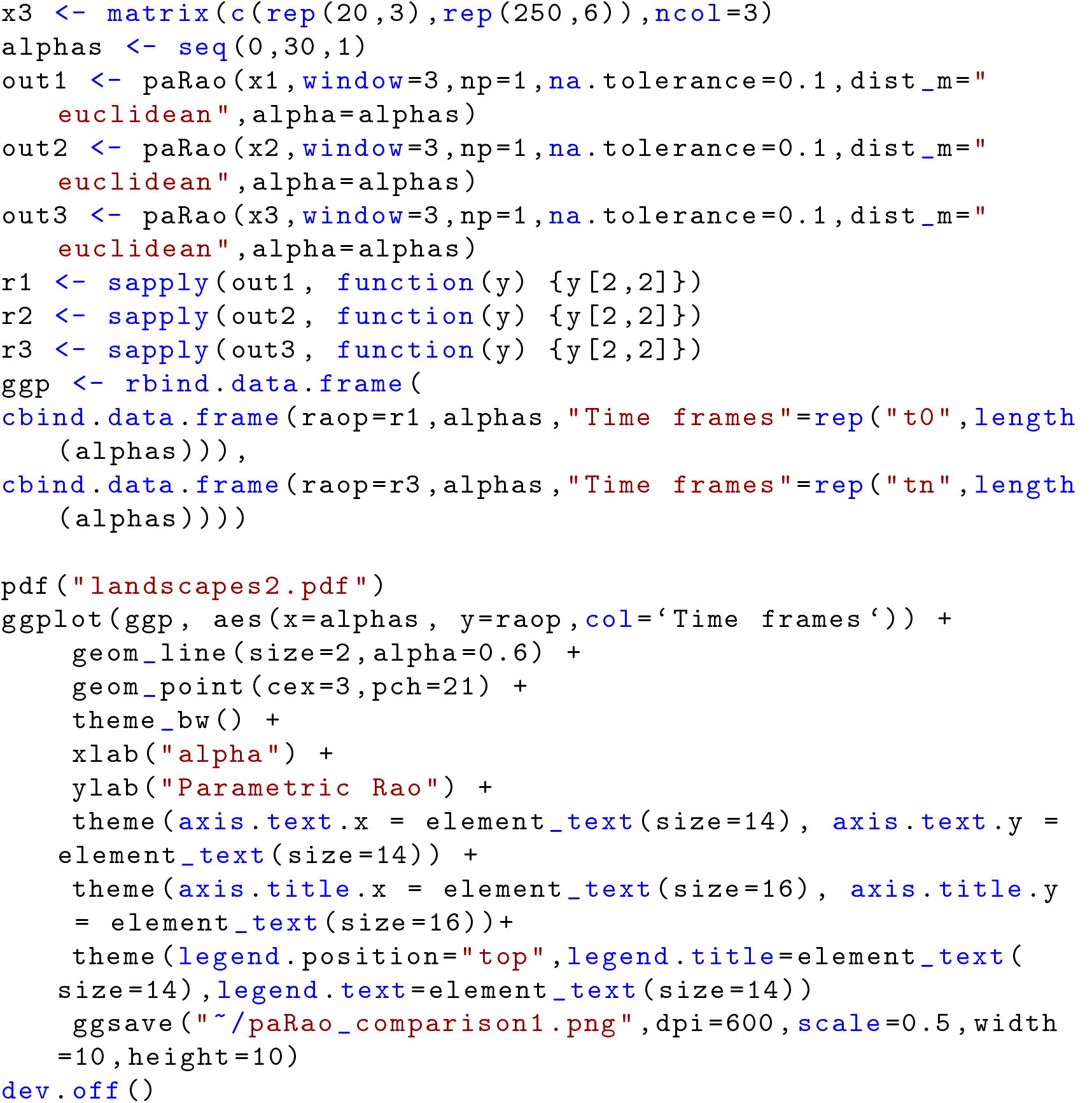

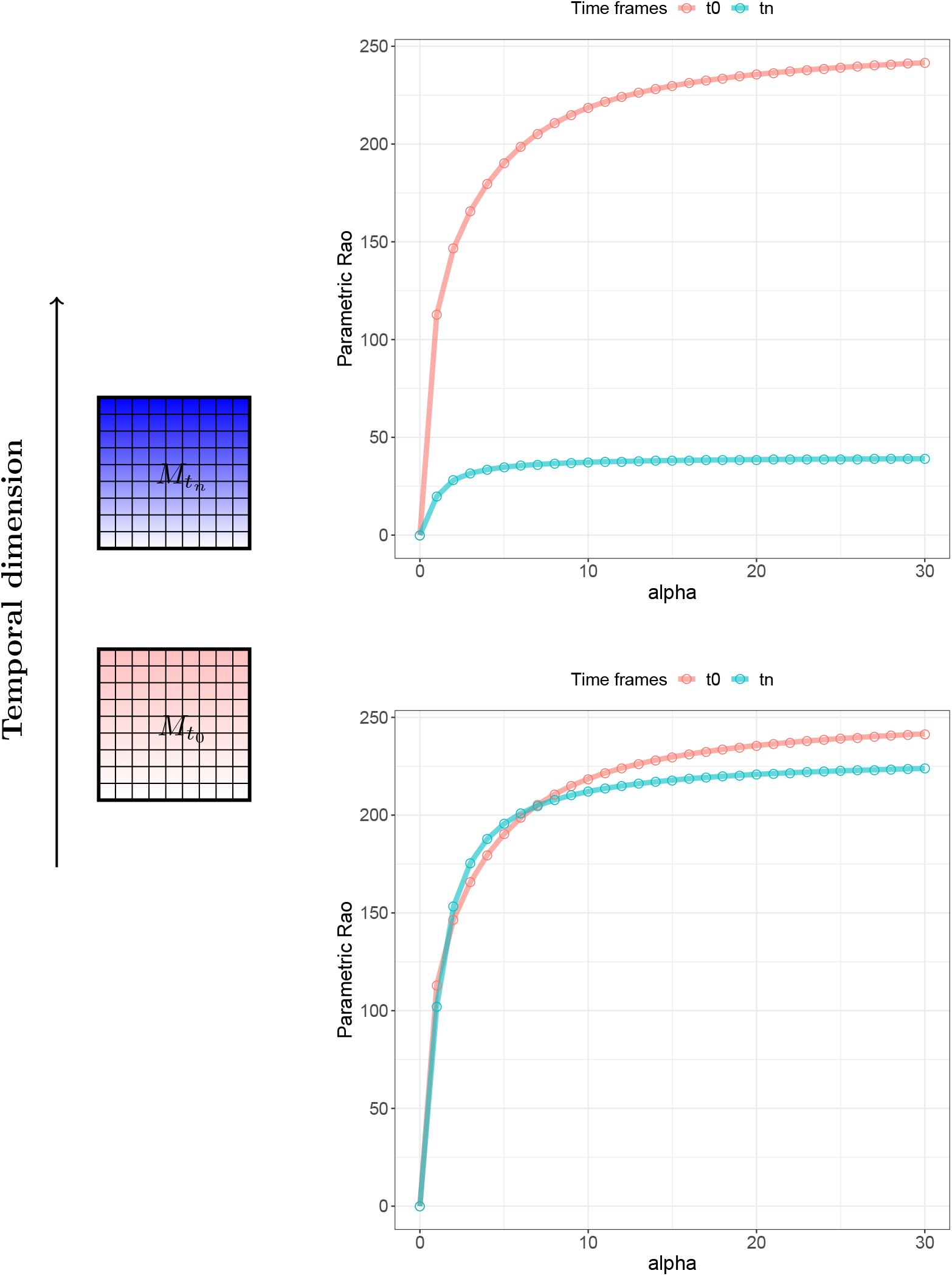

## Notes

https://cran.r-project.org/web/packages/rasterdiv/index.html

### Competing Interest Statement

The authors have declared no competing interest.

